## Supplementary Material for "Cancer cell sedimentation in 3D cultures reveals active migration regulated by self-generated gradients and adhesion sites"

#### S.1 Experiments

Overall, we performed 5 different experiments that enabled us to understand the behaviour of TNBC cells of the MDA-MB-231 cell line in 3D cultures; (i) a 3D culture experiment (control) where the cells migrated towards the glass bottom of the plate, (ii) a 3D culture experiment with administration of the drug Paclitaxel, which inhibits migration, (iii) a 3D culture RNA-sequencing experiment under Control (no treatment), and Paclitaxel treatment conditions, (iv) a 3D culture experiment that included agarose coating of the glass bottom, and (v) a 3D co-culture experiment with TNBC cells and fibroblasts.

As described in the manuscript, sedimentation was inhibited in the presence of Paclitaxel, for the three highest concentrations (0.5 μM, 0.05 μM, and 0.005 μM). This is depicted in Fig. S.1, S.2. Further analysis of bulk RNA-seq data between non-treatment and Paclitaxel-treatment conditions revealed active migration mechanisms upregulated in the non-treatment data. For the RNA-seq experiment the analysis pipeline is presented in Fig. S.3. The Gene Ontology over-representation test (Fig. S.5) revealed that many of the over-represented ontologies are linked to cellular mechanisms of migration. In particular, the Adherens Junction (AJ) signalling together with MAPK, and TGF-β have been shown to contribute to collective migration, as described in [1]. We also found that genes involved in the EMT migration mechanism did not exhibit distinct patterns of expression between non-treatment and treatment conditions (Fig. S.4). Hence, migration via the EMT mechanism was less likely to occur compared to collective migration via MAPK, TGF-β, and AJ signalling.

For the 3D culture experiment that included agarose coating of the glass bottom, as described in the manuscript, the cells in the coated sample stopped moving towards the bottom, indicating the important role of the glass in cell migration. Additionally, we discussed that the viability of the cells was reduced in the coated samples, as shown in Fig. S.6.

#### S.2 Hybrid model

As described in the manuscript, the hybrid mathematical model consisted of two parts; a continuum, and a discrete. The continuum model is a system of spatiotemporal Keller-Segel type equations that describe the spatiotemporal evolution of cancer cells and chemotactic signals. The model is then converted to a cellular automaton that is presented in Fig. S.7.

#### S.3 Cell viability assay

As described in [2], we performed a cell viability test on the 3D cultures using flow cytometry. The number of dead cells was measured for days 11, 12, and 13 after the time of seeding. For each sample, the following protocol was implemented.

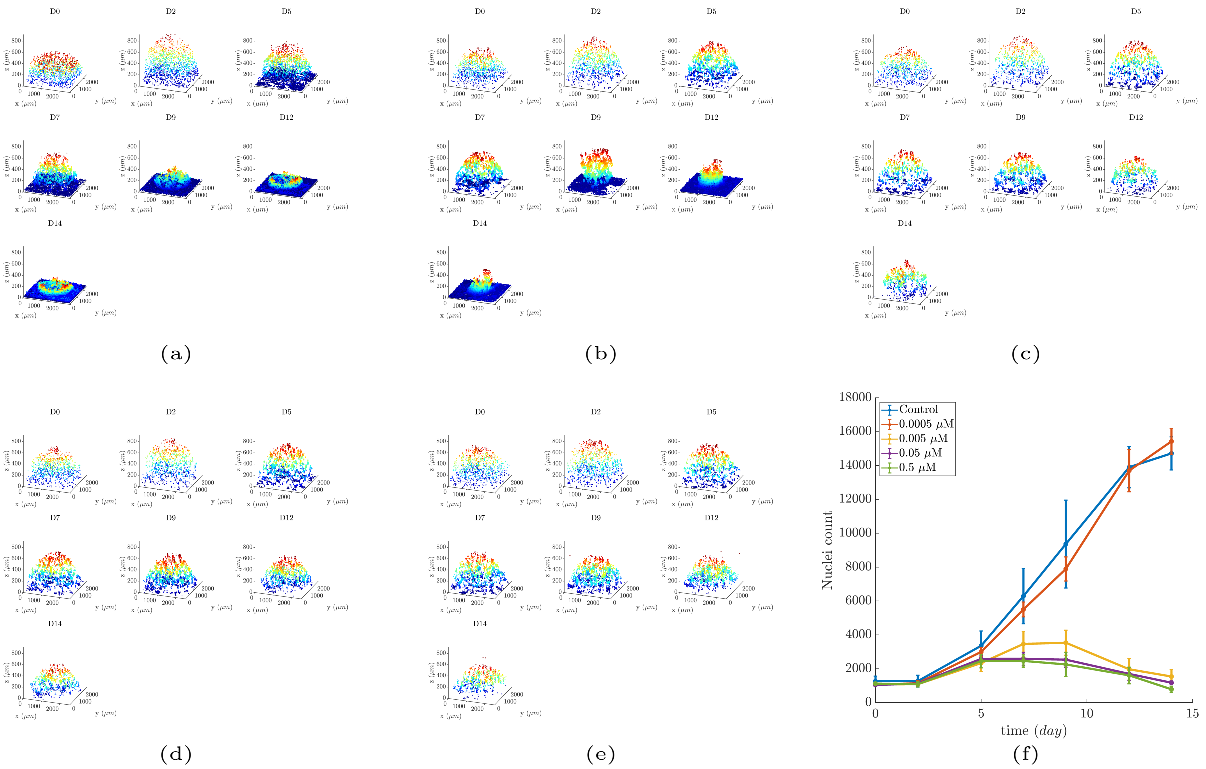


Figure S.1: Cell locations in the 3D space for (a) non-treatment (control), (b) 0.0005 μM, (c) 0.005 μM, (d) 0.05 μM, and (e) 0.5 μM of Paclitaxel, respectively. (f) Number of segmented nuclei with respect to time. We observe that the three highest doses of Paclitaxel inhibited cell movement, with the total number of cells remaining relatively constant across time, with a reduction occurring after the 9th day.

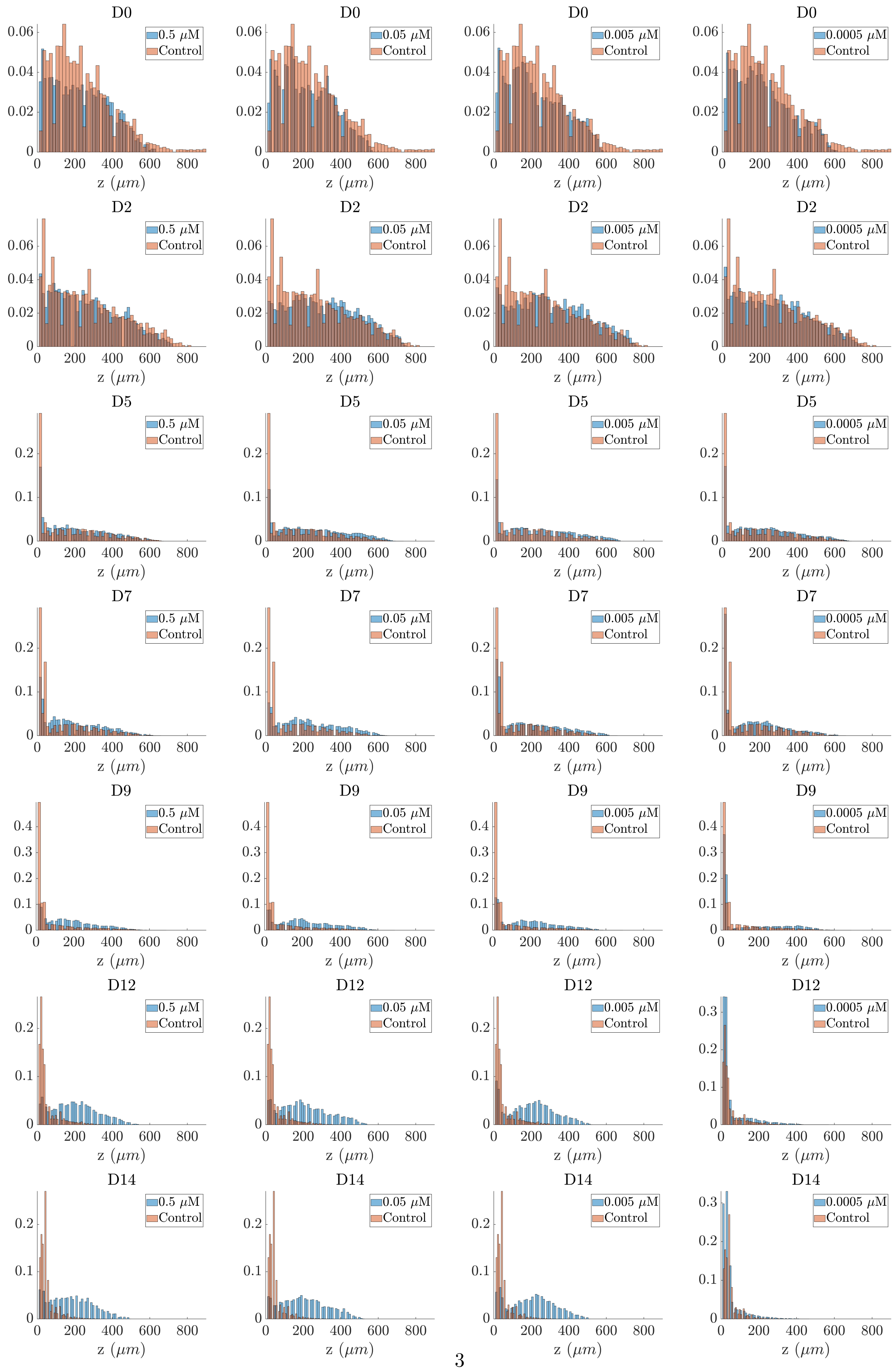




Figure S.2: Histograms of cell numbers as a function of cell culture height for control and Paclitaxel treatment.

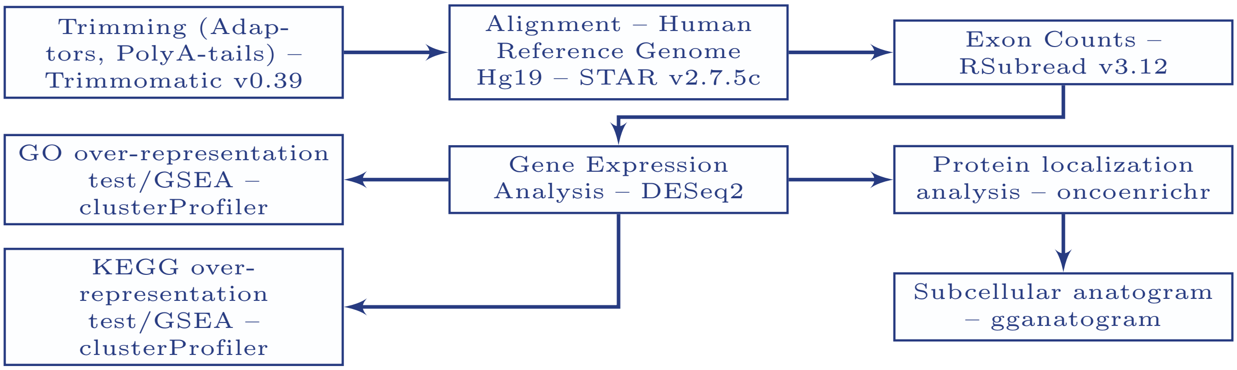


Figure S.3: RNA-seq data analysis pipeline. The differential gene expression analysis was performed between non-treatment and Paclitaxel treatment conditions. The packages used for each step are presented.

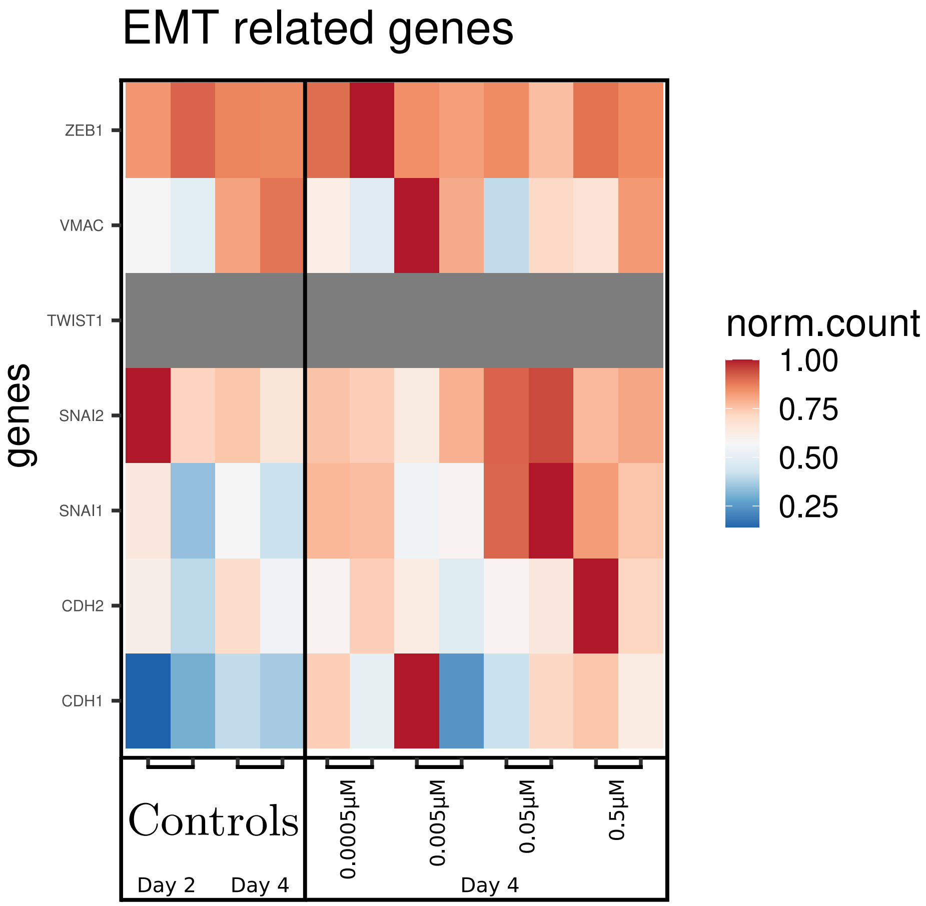


Figure S.4: The EMT signalling pathway did not exhibit a distinct pattern between treatment and non-treatment. As a result, the EMT mechanism was not considered as a plausible mechanism for migration.

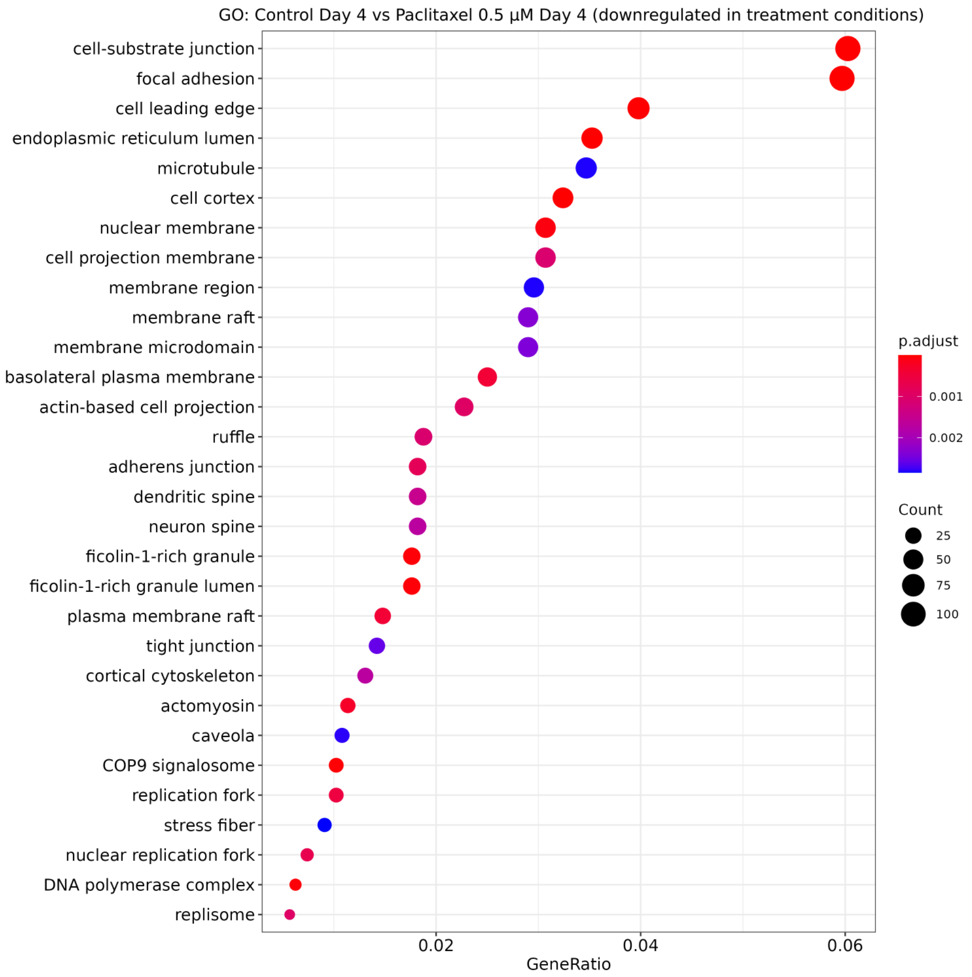


Figure S.5: Gene Ontology (GO) over-representation test. Among other ontologies related to cell movement, the Adherens Junction (AJ) is over-represented with an adjusted p-value ≈ 0.0015.

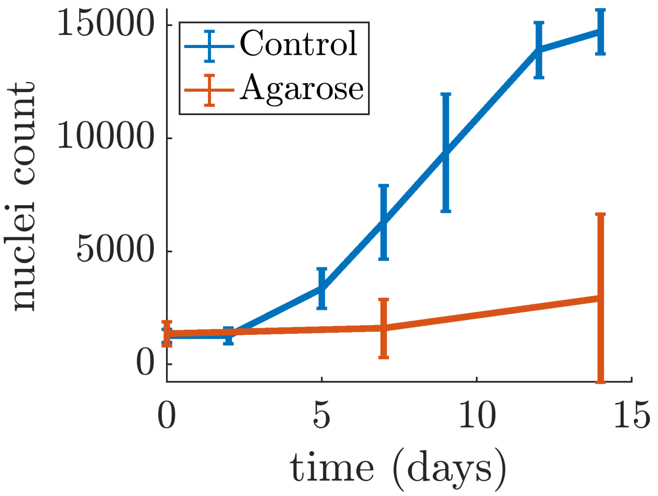


Figure S.6: Agarose coating experiment. Cell viability was reduced compared to the control, non-coating experiment. Thus, surfaces of adhesion play a major part in cell viability in *in-vitro* conditions.

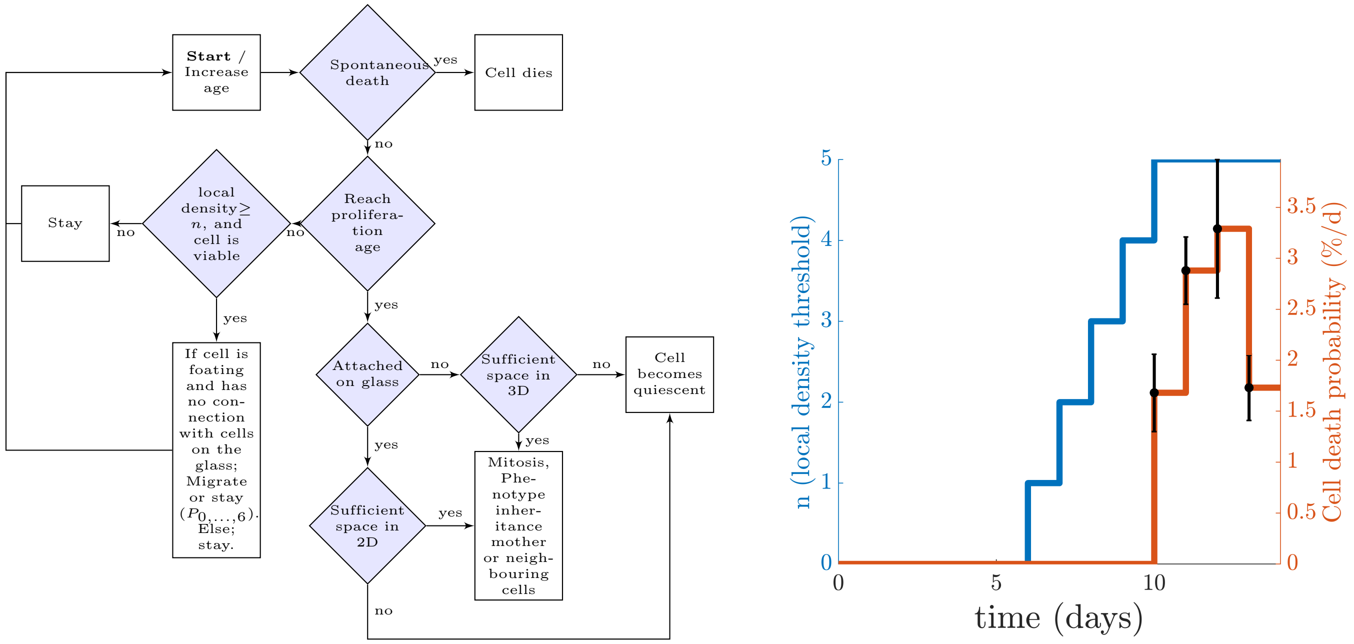


(a) (b)

Figure S.7: Cellular automaton (CA). (a) Flowchart of the CA. (b) Migration and spontaneous death probability parameters of the CA with respect to time. The migration of cells depends on the number of neighbours. The increase of the migration parameter with respect to time is based on observations made on the experiments showing that cells tended to form clusters during the later time-points. The spontaneous death probability was measured using a flow cytometry experiment.

Viability measurements for the first time-point (Day 0) was acquired from cells cultured as monolayers. Measurements obtained from every other time point correspond to cells extracted from Matrigel domes. Fresh cell culture media (DMEM (Gibco) at pH 7.2 supplemented with 10% fetal bovine serum (Wisent Bioproducts), 100 U/mL penicillin, 100 µg/mL streptomycin, and 0.25 µg/mL amphotericin B (Sigma)) was warmed at 37°C and supplemented with collagenase and dispase 1X (Sigma Aldrich, Cat. # 11097113001). Micropipettes were used to break down the cell laden Matrigel (Corning) constructs by suction force, and these were incubated for 30 minutes inside a cell culture incubator (95% relative humidity, 37°C, and 5% CO2). Cells were centrifuged at 500g for 5 minutes, the supernatant was discarded and a 0.05% trypsin (Wisent, Cat. # 325-042-CL) was used to break any cell-to-cell adhesions for a period of 5 min. Trypsin was neutralized with cell culture medium, and the cell pellet was recollected after 5 min of centrifugation at 500g. A solution containing warm (37°C) 1X DPBS and 1X propidium iodine (miltenyi biotech, Cat. # 130-093-233) was used to resuspend and incubate the collected cells for 15 minutes. Finally, cells were centrifuged again at 500g for 5 minutes, the supernatant was discarded and they were resuspended in DPBS prior to flow cytometry experimentation. All time-points were acquired in triplicates and a PI+ control was implemented using fixed (4% PFA, 5 min) and pierced cells (0.01% SDS, 1 min). Dead cells were identified using the gating strategy described in Crowley et al. [3]. The percentage of dead cells was used to directly obtain the cell death probability rate, which is presented in Fig. 4b of the manuscript.

### S.4 Continuum model calibration

The parameters of the continuum model were estimated using Bayesian inference and the Transitional Markov Chain Monte Carlo algorithm (TMCMC) [4] as implemented in the Pi4U package [5]. The TMCMC algorithm estimated posterior distributions of the model parameters given the experimental data. The process was repeated for each dataset separately across the non-treatment and treatment conditions. In total, 32 estimations were performed, and each of them evaluated 14000-20000 sets of model parameters.

The TMCMC algorithm implemented a series of resampling stages. Initially, samples from a uniform prior distribution were drawn. Then, in each stage, samples of the model parameters were drawn with respect to PDF of the corresponding stage. The plausibility weights between the proposed samples and the sample of the previous stage were computed as shown in Eq. (1). The algorithm resampled the posteriors between different stages based on the normalized weights (Eq. (2)). At the final stage, the algorithm generated samples based on the previous stage, which are distributed as follows: Eq. (3).

$$w(\underline{\theta}_{j,k}) = \frac{p_{j+1}(\underline{\theta}_{j,k})}{p_j(\underline{\theta}_{j,k})} = \frac{p(D|\underline{\theta}_{j,k}, M)^{\rho_{j+1}} p(\underline{\theta}_{j,k}|M)}{p(D|\underline{\theta}_{j,k}, M)^{\rho_j} p(\underline{\theta}_{j,k}|M)} = p(D|\underline{\theta}_{j,k}, M)^{\rho_{j+1}-\rho_j} \tag{1}$$

$$\underline{\tilde{w}}_{j,l} = \frac{w(\underline{\theta}_{j,l})}{\sum_{l=1}^{N_j} w(\underline{\theta}_{j,l})} \tag{2}$$

$$p(\underline{\theta}) \sim p(D|\underline{\theta}, M)p(\underline{\theta}|M) \tag{3}$$

The average and standard deviation values of the parameters, as calculated from their posterior distributions across the examined conditions are given in Tables S.1-S.5. The posterior distributions of the model parameters for representative datasets across each condition are depicted in Fig. S.8.

### S.5 Normalized Root Mean Squared Error

The differences between the *in-vitro* and *in-silico* estimated cell density profiles were calculated using the Normalized Root Mean Squared Error (NRMSE) as defined in (4), where $u_e$, $u_s$ are the arrays containing the experimental and simulated density profiles, respectively.

$$\text{NRMSE} = \frac{1}{\max(u_e) - \min(u_e)} \sqrt{\frac{\sum_{i=1}^{N} (u_s^i - u_e^i)^2}{N}} \tag{4}$$

| Dataset | $D_u \in [10^{-3}, 2] \times 10^{-3}$ $(mm^2d^{-1})$ | $s \in [1.8, 3] \times 10^{-1}(d^{-1})$ | $\chi \in [0.8, 5] \times 10^{-2}(mm^2d^{-1})$ | $D_f \in [10^{-3}, 2] \times 10^{-3}$ $(mm^2d^{-1})$ |
|---|---|---|---|---|
| 1 | $1.999 \pm 0.001$ | $1.843 \pm 0.001$ | $0.833 \pm 0.001$ | $1.992 \pm 0.003$ |
| 2 | $1.698 \pm 0.138$ | $2.164 \pm 0.140$ | $0.819 \pm 0.014$ | $1.792 \pm 0.120$ |
| 3 | $1.872 \pm 0.070$ | $1.805 \pm 0.003$ | $0.883 \pm 0.028$ | $1.724 \pm 0.149$ |
| 4 | $0.006 \pm 0.001$ | $1.841 \pm 0.017$ | $0.830 \pm 0.028$ | $1.404 \pm 0.050$ |
| 5 | $1.995 \pm 0.004$ | $1.806 \pm 0.005$ | $0.802 \pm 0.001$ | $1.983 \pm 0.006$ |
| 6 | $1.807 \pm 0.001$ | $2.463 \pm 0.001$ | $0.936 \pm 0.001$ | $1.229 \pm 0.001$ |
| 7 | $0.205 \pm 0.014$ | $1.803 \pm 0.002$ | $1.063 \pm 0.068$ | $0.895 \pm 0.108$ |
| 8 | $0.328 \pm 0.026$ | $1.998 \pm 0.016$ | $2.050 \pm 0.069$ | $1.748 \pm 0.019$ |
| 9 | $0.244 \pm 0.022$ | $1.823 \pm 0.011$ | $2.414 \pm 0.166$ | $1.703 \pm 0.124$ |
| 10 | $1.854 \pm 0.085$ | $2.243 \pm 0.020$ | $0.803 \pm 0.002$ | $0.118 \pm 0.019$ |
| 11 | $1.951 \pm 0.035$ | $2.350 \pm 0.073$ | $0.804 \pm 0.003$ | $1.840 \pm 0.034$ |
| 12 | $1.514 \pm 0.090$ | $1.871 \pm 0.036$ | $0.874 \pm 0.024$ | $0.032 \pm 0.003$ |

Table S.1: Average and standard deviation of the inferred parameter values of the continuum KS model across all datasets for the non-treatment condition (control). The prior PDF boundaries used for the parameter estimation are shown in the header of the table.

| Dataset | $D_u \in [0, 2] \times 10^{-3}$ $(mm^2d^{-1})$ | $s \in [1.92, 2.08] \times 10^{-1}(d^{-1})$ | $k \in [0.0, 0.2](d^{-1})$ | $\chi \in [0, 5] \times 10^{-2}(mm^2d^{-1})$ | $D_f \in [1.2, 1.5] \times 10^{-3}$ $(mm^2d^{-1})$ |
|---|---|---|---|---|---|
| 1 | $1.5510 \pm 0.0689$ | $2.0417 \pm 0.0041$ | $0.0032 \pm 0.0016$ | $1.0475 \pm 0.0188$ | $1.2866 \pm 0.0083$ |
| 2 | $1.2040 \pm 0.0088$ | $2.0350 \pm 0.0007$ | $0.0506 \pm 0.0074$ | $0.0106 \pm 0.0053$ | $1.2418 \pm 0.0003$ |
| 3 | $1.7316 \pm 0.0246$ | $2.0810 \pm 0.0032$ | $0.0028 \pm 0.0029$ | $1.1428 \pm 0.0096$ | $1.4006 \pm 0.0116$ |
| 4 | $1.1329 \pm 0.0005$ | $2.0840 \pm 0.0000$ | $0.0100 \pm 0.0003$ | $2.0913 \pm 0.0004$ | $1.2693 \pm 0.0001$ |
| 5 | $1.7862 \pm 0.1308$ | $2.0779 \pm 0.0021$ | $0.0520 \pm 0.0092$ | $1.9857 \pm 0.0672$ | $1.2636 \pm 0.0043$ |

Table S.2: Average and standard deviation of the inferred parameter values of the continuum KS model across all datasets for Paclitaxel treatment with concentration 0.0005 µM. The prior PDF boundaries used for the parameter estimation are shown in the header of the table.

| Dataset | $D_u \in [0.0, 10^{-2}] \times 10^{-2}$ $(mm^2d^{-1})$ | $s \in [1.92, 2.08] \times 10^{-1}(d^{-1})$ | $k \in [2, 5] \times 10^{-1}(d^{-1})$ | $\chi \in [0.0, 10^{-3}] \times 10^{-2}(mm^2d^{-1})$ | $D_f \in [1.2, 1.5] \times 10^{-3}$ $(mm^2d^{-1})$ |
|---|---|---|---|---|---|
| 1 | $0.0553 \pm 0.0049$ | $2.0638 \pm 0.0083$ | $2.0081 \pm 0.0056$ | $0.0004 \pm 0.0001$ | $1.2559 \pm 0.0120$ |
| 2 | $0.0675 \pm 0.0017$ | $2.0822 \pm 0.0008$ | $2.0009 \pm 0.0006$ | $0.0004 \pm 0.0001$ | $1.2439 \pm 0.0029$ |
| 3 | $0.0755 \pm 0.0041$ | $2.0822 \pm 0.0010$ | $2.0005 \pm 0.0005$ | $0.0005 \pm 0.0001$ | $1.2719 \pm 0.0068$ |
| 4 | $0.0688 \pm 0.0011$ | $2.0826 \pm 0.0008$ | $2.0015 \pm 0.0010$ | $0.0004 \pm 0.0001$ | $1.2448 \pm 0.0023$ |
| 5 | $0.0757 \pm 0.0053$ | $2.0801 \pm 0.0030$ | $2.0028 \pm 0.0028$ | $0.0003 \pm 0.0001$ | $1.3129 \pm 0.0114$ |

Table S.3: Average and standard deviation of the inferred parameter values of the continuum KS model across all datasets for Paclitaxel treatment with concentration 0.005 µM. The prior PDF boundaries used for the parameter estimation are shown in the header of the table.

| Dataset | $D_u \in [0.0, 10^{-2}] \times 10^{-3}$ $(mm^2d^{-1})$ | $s \in [1.92, 2.08] \times 10^{-1}(d^{-1})$ | $k \in [2, 5] \times 10^{-1}(d^{-1})$ | $\chi \in [0.0, 10^{-3}] \times 10^{-2}(mm^2d^{-1})$ | $D_f \in [1.2, 1.5] \times 10^{-3}$ $(mm^2d^{-1})$ |
|---|---|---|---|---|---|
| 1 | $0.0067 \pm 0.0001$ | $2.0836 \pm 0.0003$ | $2.0009 \pm 0.0006$ | $0.0005 \pm 0.0001$ | $1.2560 \pm 0.0054$ |
| 2 | $0.0054 \pm 0.0003$ | $2.0810 \pm 0.0013$ | $2.0023 \pm 0.0016$ | $0.0004 \pm 0.0001$ | $1.2745 \pm 0.0086$ |
| 3 | $0.0072 \pm 0.0004$ | $2.0509 \pm 0.0078$ | $2.0105 \pm 0.0091$ | $0.0002 \pm 0.0001$ | $1.3234 \pm 0.0254$ |
| 4 | $0.0072 \pm 0.0007$ | $2.0803 \pm 0.0026$ | $2.0022 \pm 0.0018$ | $0.0006 \pm 0.0001$ | $1.3097 \pm 0.0303$ |
| 5 | $0.0053 \pm 0.0006$ | $2.0768 \pm 0.0061$ | $2.0054 \pm 0.0053$ | $0.0007 \pm 0.0002$ | $1.3945 \pm 0.0460$ |

Table S.4: Average and standard deviation of the inferred parameter values of the continuum KS model across all datasets for Paclitaxel treatment with concentration 0.05 µM. The prior PDF boundaries used for the parameter estimation are shown in the header of the table.

| Dataset | $D_u \in [0.0, 10^{-2}] \times 10^{-3}$ $(mm^2d^{-1})$ | $s \in [1.92, 2.08] \times 10^{-1}(d^{-1})$ | $k \in [2, 5] \times 10^{-1}(d^{-1})$ | $\chi \in [0.0, 10^{-3}] \times 10^{-2}(mm^2d^{-1})$ | $D_f \in [1.2, 1.5] \times 10^{-3}$ $(mm^2d^{-1})$ |
|---|---|---|---|---|---|
| 1 | $0.0097 \pm 0.0003$ | $2.0030 \pm 0.0127$ | $2.5438 \pm 0.0211$ | $0.0003 \pm 0.0001$ | $1.2738 \pm 0.0261$ |
| 2 | $0.0081 \pm 0.0001$ | $2.0836 \pm 0.0003$ | $2.5327 \pm 0.0137$ | $0.0001 \pm 0.0001$ | $1.2734 \pm 0.0041$ |
| 3 | $0.0054 \pm 0.0003$ | $2.0815 \pm 0.0012$ | $2.0015 \pm 0.0013$ | $0.0007 \pm 0.0001$ | $1.3193 \pm 0.0189$ |
| 4 | $0.0067 \pm 0.0007$ | $2.0682 \pm 0.0068$ | $2.0078 \pm 0.0066$ | $0.0005 \pm 0.0001$ | $1.3828 \pm 0.0221$ |
| 5 | $0.0072 \pm 0.0007$ | $2.0701 \pm 0.0056$ | $2.0070 \pm 0.0053$ | $0.0004 \pm 0.0001$ | $1.2670 \pm 0.0212$ |

Table S.5: Average and standard deviation of the inferred parameter values of the continuum KS model across all datasets for Paclitaxel treatment with concentration 0.5 µM. The prior PDF boundaries used for the parameter estimation are shown in the header of the table.

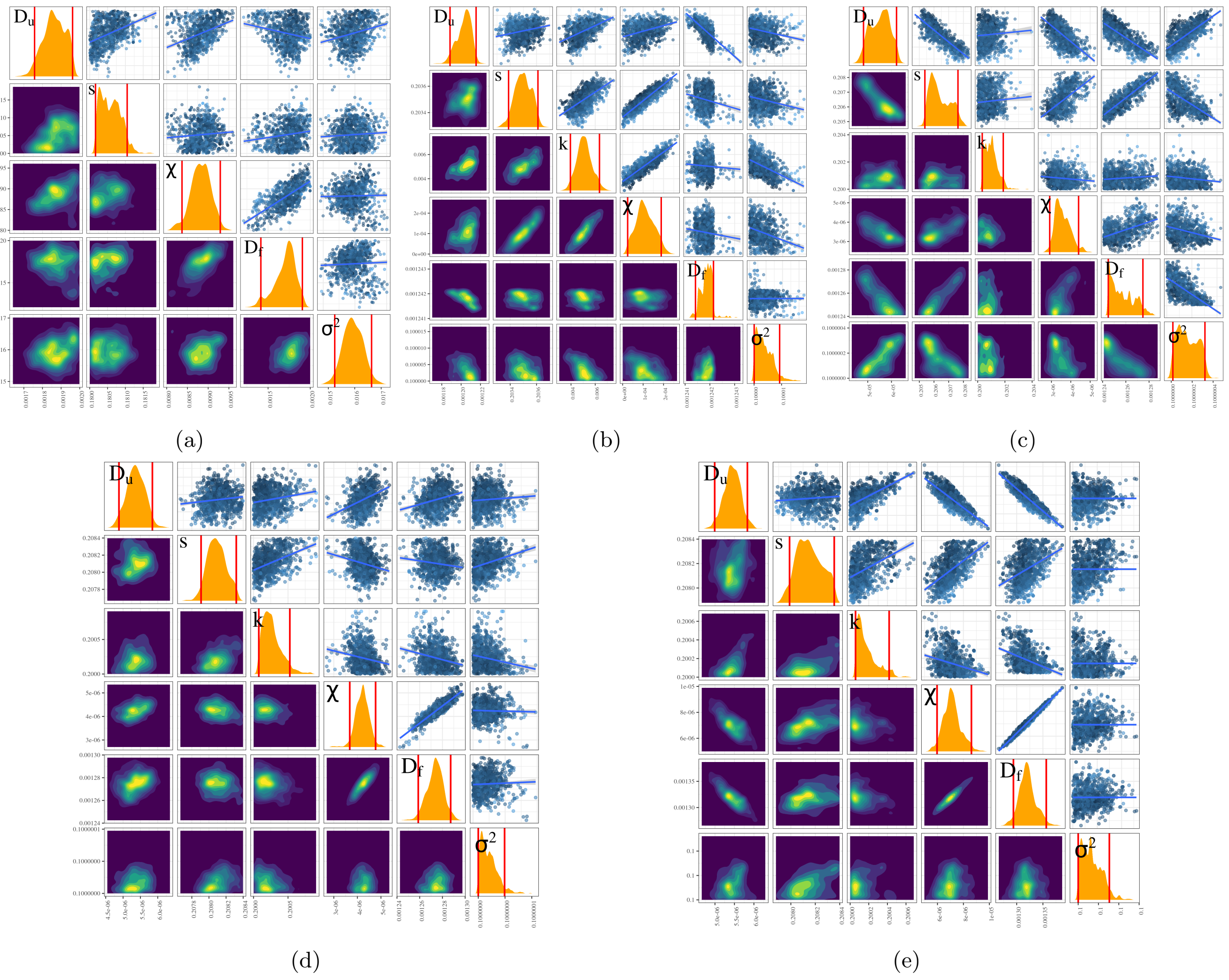

Figure S.8: Model calibration. Posterior distributions of the model parameters for representative datasets corresponding to (a) non-treatment conditions, and (b) 0.0005 µM, (c) 0.005 µM, (d) 0.05 µM, (e) 0.5 µM of Paclitaxel treatment. Upper diagonal: Projected TMCMC samples of the posterior distribution in 2D space. Diagonal: Marginals of the joint posterior obtained via kernel densities. Red lines denote the 95% credible intervals. Lower diagonal: 2D projected densities of the posterior distribution obtained using 2D kernel densities.

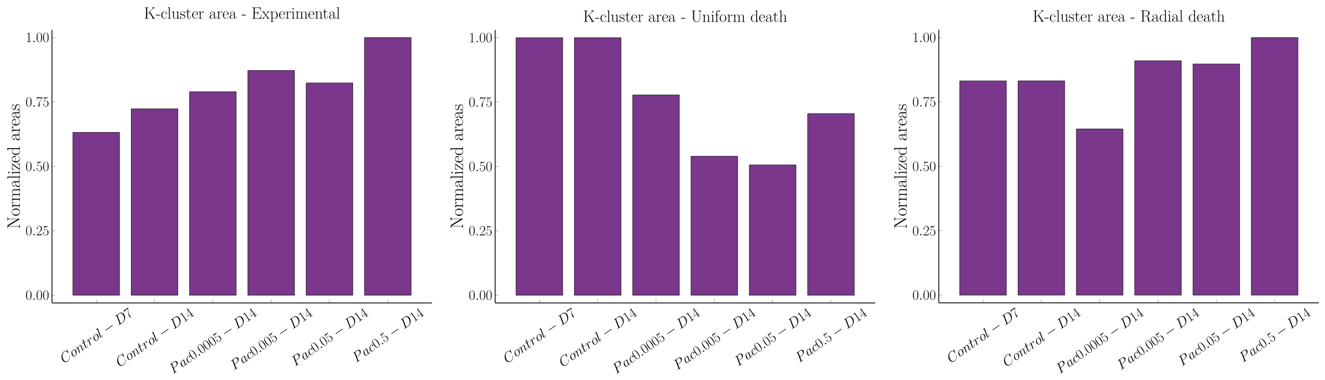


(a) (b) (c)

Figure S.9: Clustering abundance for (a) Experiments, (b) Simulations with uniform death probability across the space, and (c) Simulations with a radial decrease in cell death probability towards the center of the space.

### S.6 Model validation analysis

To perform model validation, we implemented the Complete Spatial Randomness test using Ripley's K-function, and we calculated the Nearest-Neighbour (NN), and the Inter-Nucleic (IN) Euclidean distance distributions. The results obtained from the hybrid model were compared with the results obtained from the experimental data, suggesting good agreement overall. In this section, we present the results obtained from the calculation of the IN distances between the cells (Fig. S.10). The distributions remained relatively stable across time and condition, indicating that the behaviour of cells was the same across the examined space.

The CSR test revealed pronounced clustered patterns across all datasets, with the experimental data exhibiting more pronounced clustering patterns for increasing time and dose (Fig. S.9a). The degree of clustering was quantified by calculating the area under the curve of the K-function ($\text{AUC}_K$). The normalized $\text{AUC}_K$, which is defined as, $\hat{\text{AUC}}_{K,i} = \text{AUC}_{K,i}/\max(\text{AUC}_K)$. More pronounced clustering patterns had increased values of $\hat{\text{AUC}}_K$ as shown in Fig. S.9a. In contrast to the experimental data, the simulation results yielded less pronounced clustering patterns for higher drug doses (Fig. S.9b). We attributed this result to the fact that the *in-silico* cells had a uniform probability of death across the space, while in the experiment, the drug concentration may have formed a gradient decreasing towards the core of the scaffold. To examine this hypothesis, we repeated the simulations, this time introducing a gradient to the cell death probability that decreased radially towards the core of the scaffold. In turn, the death probability of a cell found in $(x_i, y_i, z_i)$ became,

$$d_{r,i} = p_k \frac{\sqrt{(x_i - x_c)^2 + (y_i - y_c)^2 + (z_i - z_c)^2}}{0.5\sqrt{x_c^2 + y_c^2 + z_c^2}} \tag{5}$$

where $p_k = k\mathrm{d}t$ is the death probability determined by the death rate, $k$, of the continuum model, and $(x_c, y_c, z_c)$ the center of the space. The radius dependent death probability yielded more similar results to the experiment, i.e. a clustering index that increased as a function of drug dose (Fig. S.9c).

As mentioned in the Discussion section C of the manuscript, the mechanism behind the increasing clustering with respect to time is the advection that biases the movement of the cells towards the increase of signal concentration. On the other hand, the diffusion mechanism would likely result in less pronounced clustering. To demonstrate the net effect of advection together with diffusion we performed 2 additional simulations, one with low advection constants and high diffusion constants, as well as another with high advection and low diffusion constants, respectively. In total, 12 simulations for each of the 2 cases were performed. Subsequently, we used the resulted cell distributions to perform the CSR test. Finally, we calculated the are under the curve of the K-function ($\text{AUC}_K$), and we compared it to the values of the diffusion and advection constants. The results presented in Fig. S.11 show that the combination of low advection and high diffusion constants produced considerably lower $\text{AUC}_K$ values than those with high advection and low diffusion constants. The statistical significance of the resulted $\text{AUC}_K$ between these two cases was calculated using the Kruskal-Wallis paired test,

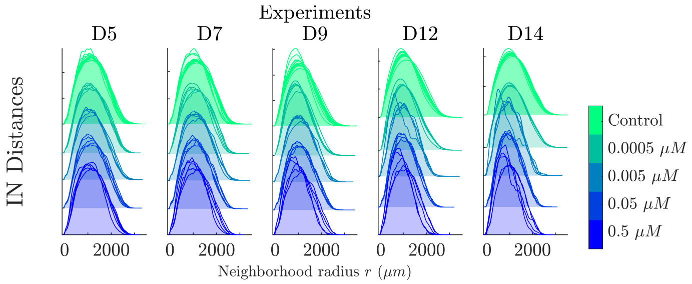


(a)

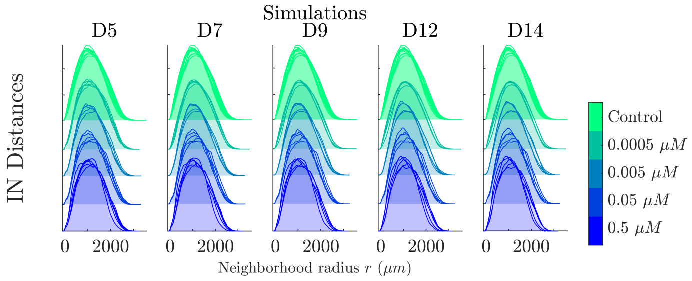


(b)

Figure S.10: Inter-Nucleic Euclidean distance distributions between (a) experiments, and (b) simulations across time. Overall, the distributions remained stable across time and treatment condition in both experiments and simulations.

and the p-value was found to be $5.4 \times 10^{-5}$.

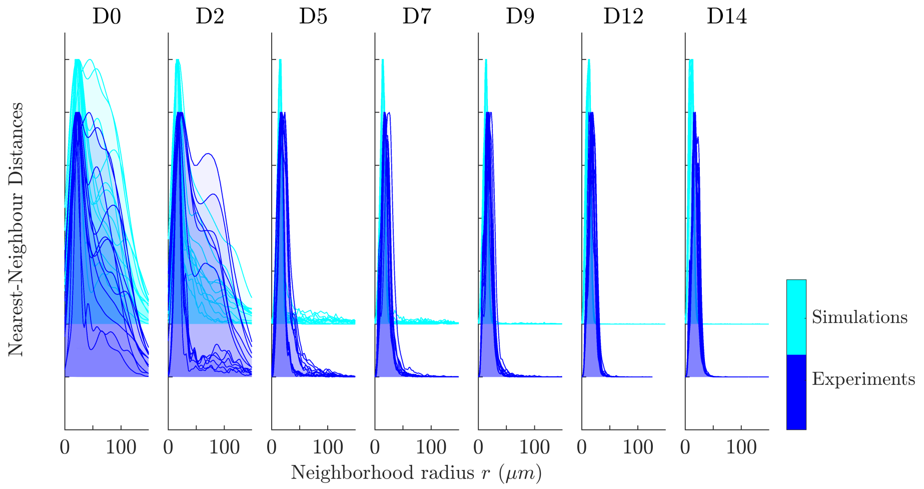


Figure S.11: Nearest-Neighbour Euclidean distance distributions for the non-treatment conditions across time. The NN distances initially formed wide distributions that gradually became narrower around lower neighbourhood radii values with respect to time, across all samples, with similar characteristic peaks at ∼15 µm.

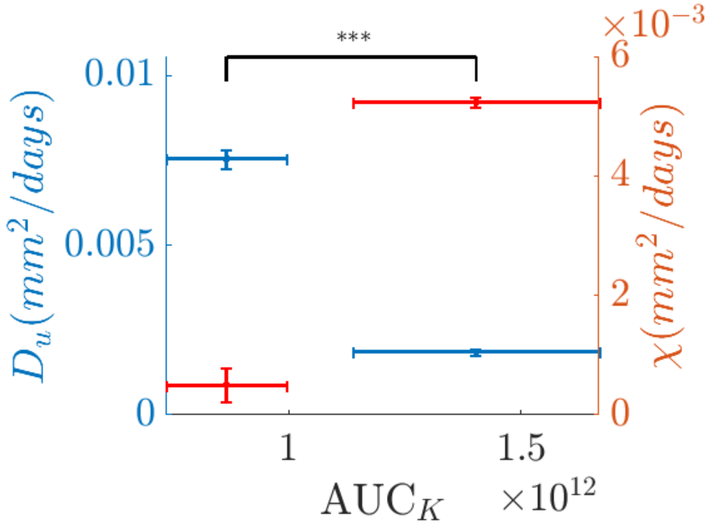


Figure S.12: Relationship between diffusion and advection mechanisms with morphological patterns. The K-function was calculated for 12 simulations corresponding to the following two cases: one with low advection and high diffusion constants, as well as another with high advection and low diffusion constants. The area under the resulted K-function ($\text{AUC}_K$) was calculated, revealing that the combination low advection and high diffusion constants yielded a considerably lower $\text{AUC}_K$ than the combination of high advection and low diffusion constants. The statistical significance of this result was calculated using the Kruskal-Wallis test, and it was found that p-value$\approx 5.4 \times 10^{-5}$.
